# Interaction of Gut-Microbial Amyloids with Endogenous Amyloids can Drives Microglial Hyperactivity and Neuroinflammation in Alzheimer’s Disease

**DOI:** 10.1101/2025.01.15.633290

**Authors:** Helen Forgham, Eduardo A. Albornoz, Giovanni Pietrogrande, Jiayuan Zhu, Ka Hang Karen Chung, Muhammad S. Tahir, Syed Aoun Ali, Ruirui Qiao, Aleksandr Kakinen, Ernst J. Wolvetang, Trent M. Woodruff, Julio Aguado, Daniel E. Otzen, Thomas P. Davis, Ibrahim Javed

## Abstract

Gut bacteria have emerged as silent drivers in the pathology of Alzheimer’s disease (AD). They also make amyloids with structure analogue to pathological amyloids and have potential to cross-seed and propagate in a prion-like manner. AD is characterised by the accumulation of mature extracellular Amyloid-β (Aβ) plaques which are surrounded by inflammatory microglia. We report that exposure to interspecies microbial amyloids of FapC (fimbriae) and CsgA (curli) from opportunistic gut pathogens *Pseudomonas aeruginosa* and *Escherichia coli* hyperactivates microglia against Aβ fibrils. Microbial amyloids and Aβ fibrils converge in phagocytic compartments through subsequent internalization, not observed with Aβ fibrils alone. This convergence promotes pro-inflammatory microglia with a defective proteome similar to those observed in AD brains. The resulting clusters develop a pro-inflammatory, indigestible interactome that is eventually regurgitated, inducing progressive degeneration in bystander neurons and ultimately leading to cognitive decline. Collectively, these findings provide compelling evidence that microbial amyloids can trigger progressive AD pathology through microglia-driven neuroinflammation.

## Introduction

The gut microbiome influences microglia maturation and has recently been recognised as central to Alzheimer’s disease (AD) pathology.^1,2^ Many opportunistic gut pathogens produce scaffolding amyloids that mimic the folding complexities of human Amyloid-β (Aβ).^3^ The faecal samples from human has confirmed the presence of wide range of biofilms-associated microbial-amyloids in the gut with potential to induce neurodegeneration.^4^ However, how these microbial amyloid can access brain tissue and what is the mechanism they use to trigger or mediate pathological paradigm is unclear.^5^ Previously, they were reported to directly cross-seed the amyloidosis of Aβ and α-Synuclein (aSyn).^6,7^ Also, due to their stable cross-β architecture, microbial amyloids can access extra intestinal tissues in a prion-like manner and co-localize with pathological amyloids.^8,9^ The aging vagus nerve, gut and blood-brain barriers can also create a “leaky” window, allowing gut microbial metabolites to access the brain.^10,11^ Thus, it is reasonable to hypothesise that microbial amyloids or their fragmented seeds could access the brain and incite progressive AD neuropathology.^7,12^ However, to date, no direct evidence has been found to support the existence of this toxic trifecta - microglia, gut pathogens and Aβ – in the onset and progression of AD.

A major histopathological hallmark of Alzheimer’s disease (AD) is the accumulation of extracellular aggregated Amyloid-β (Aβ) protein plaques.^13,14^ Aβ oligomers appear early in the aggregation pathway directly triggering pro-inflammatory microglia and disrupting neuronal membrane.^15,16^ Mature Aβ fibrils have pro-inflammatory microglia localised to the plaques where they contribute to progressive inflammation and substantial neuronal loss.^17,18^ Based on the facts that microbial amyloids can cross-seed pathological amyloids, found in the gut and can access tissues outside the gut, it is reasonable to hypothesise that microbial amyloids can light-up the hyperactivity of microglia against Aβ amyloids.

FapC and CsgA proteins, secreted by opportunistic gut-pathogens of *Pseudomonas aeruginosa* and *Escherichia coli*, self-assemble into amyloid fibrils that strengthen biofilm.^3^ Here, we used a combination of cellular, organoids and zebrafish models and found that FapC and CsgA seeds act as catalysts and hyperactivate microglia against mature Aβ fibrils. Phagocytic convergence of Aβ fibrils and bacterial amyloid seeds (BAS) results in the formation of clusters within microglia. The clusters develop an inflammatory interactome that is subsequently excreted from the cell, inducing toxicity to bystander neurons and driving an inflammatory phenotype in microglia, leading to neurodegeneration.

Our results suggest a novel pathway whereby Aβ toxicity is greatly exacerbated through a microglia-mediated interaction with microbial amyloids, reconciling the mismatch between the relatively low immunogenicity of Aβ plaques *in vitro* and the strong neuroinflammatory response associated with Aβ plaques observed in postmortem samples.

## Results

### Fragmented seeds (BAS) of microbial amyloids in combination with amyloid-β (Aβ) fibrils hyperactivate microglia towards a pro-inflammatory phenotype

Bacterial amyloid seeds (BAS) were generated through the sonication of a mixture of FapC and CsgA amyloids.^7^ The rationale behind using a 1:1 mixture of FapC and CsgA was that microbial amyloids from *P. aeruginosa* and *E. coli* facilitate interspecies biofilm formation, particularly within the gut.^19,20^ We began by examining the phagocytic function of the human microglial cell line (HMC3) by exposing them to 0.1 µM (monomer units) of BAS for 24 h before imaging with phase contrast microscopy (**Fig. 1A**). Microglia displayed an accumulation of BAS, which appeared as non-uniform, irregularly shaped intracellular aggregates (**Fig. 1B**). Similar phagocytic patterns were not observed with Aβ fibrils or full-length BAS at a similar concentration, or with lipopolysaccharide (LPS) used as a positive control for microglia activation (**Fig. S1**). Phagocytic uptake of BAS stained with AF647 fluorescence dye was confirmed by Z-stacked image penetrating through the whole cell (**Vid. S1**) and was found to be non-cytotoxic to microglial cells (**Fig. 1C, D**).

**Fig. 1:**
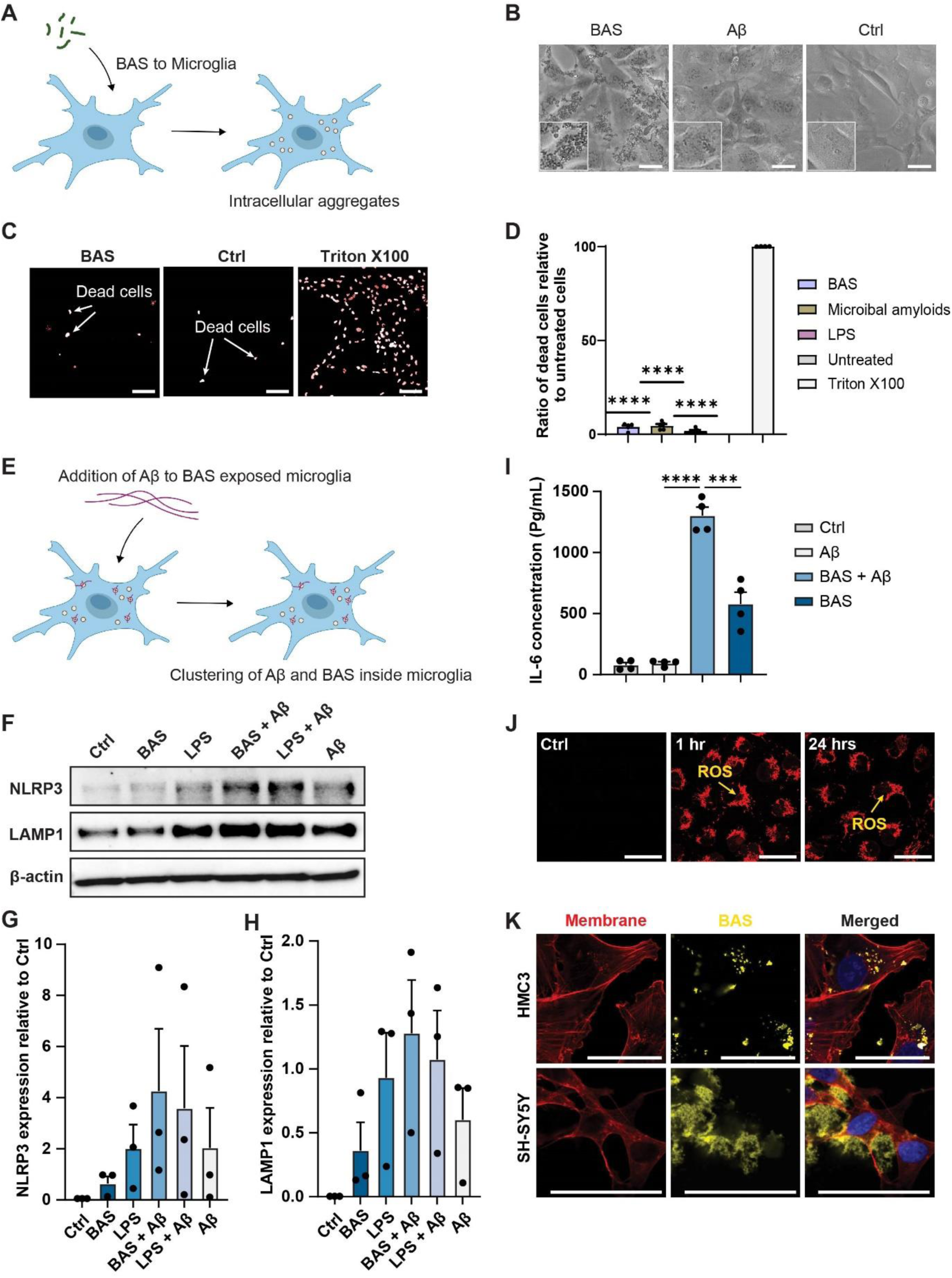
Microglia polarise to a pro-inflammatory phenotype upon exposure to BAS followed by Aβ. **A**) Illustration depicting bacterial amyloid seeds (BAS) phagocytosed by microglia leading to extensive intracellular aggregates. **B**) Phase contrast image showing intracellular aggregates are specific to BAS phagocytosis (scale bar 20 µm). **C, D**) Cytotoxicity in microglia visualised and quantified using DRAQ7 immunofluorescence (scale bar 200 µm); ****P ≤ 0.0001. **E**) Illustration demonstrating the cluster formation from convergence of BAS and Aβ species inside of phagocytic compartments. **F**) Western blotting image demonstrating increased expression of inflammasome and phagocytic markers (NLRP3 and LAMP1) relative to BAS, Aβ and lipopolysaccharide exposure. **G, H**) Graphs demonstrating semi-quantification of protein expression for NLRP3 and LAMP1, respectively. **I**) Graph illustrating IL-6 expression in media taken from HCM3 cells exposed to BAS followed by Aβ, or BAS and Aβ as lone agents. IL-6 secretion is significantly upregulated in the cells exposed to both BAS and Aβ relative to lone agents (***P ≤ 0.001, ****P ≤ 0.0001). **J**) Representative immunofluorescence image demonstrating HCM3 cells exposed to BAS followed by Aβ promotes sustained reactive oxygen species generation (red fluorescence generated upon oxidation), scale bar 50 µm. **K**) Representative immunofluorescence image demonstrating compact bundling of BAS inside phagocytic compartments of HCM3 cells. Conversely, BAS are observed as fluffy cloud like extracellular entities when exposed to SH-SY5Y neuronal cells at high relative concentration (scale bar 50 µm). All The data are representative of *n* = 3 independent experiments. Statistical values are calculated by one sample t test.

Next, we aimed to determine whether exposure to BAS alone, or in combination with Aβ, would promote the transition of microglial cells from a quiescent state to a polarised pro-inflammatory phenotype (**Fig. 1E**) using western blotting. Lysates from microglia cells previously exposed to BAS, Aβ, LPS, BAS + Aβ or LPS + Aβ were assessed for the expression of NLRP3 and LAMP1, markers for inflammasome and phagosomes, respectively. BAS alone did not significantly increase the expression of these two proteins compared to untreated cells (**Fig. 1F**). In contrast, HMC3 cells exposed to BAS + Aβ showed an increased expression of both NLRP3 and LAMP1, though this varied across three independent experiments (**Fig. 1G, H**).

Since Aβ has been reported to induce microglial dysfunction through lipid droplet formation,^21^ we ruled out the possibility that BAS could cause a similar effect by conducting Red O staining. Microglia exposed to BAS tested negative for lipid droplet formation (**Fig. S2**). We examined the secretion of cytokine IL-6 and the generation of reactive oxygen species (ROS) as proxies for a pro-inflammatory phenotype. IL-6 levels in the supernatant of microglia cells were significantly higher following exposure to BAS + Aβ compared to cells exposed to either BAS or Aβ alone (**Fig. 1I**). Furthermore, confocal live-cell microscopy and Z-stack imaging of HMC3 cells exposed to BAS + Aβ, imaged after 1 hour, showed extensive, bright cytoplasmic CellROX fluorescence (red) localized around the nuclei, with comparable intensity 24 hours after initial exposure (**Fig. 1J, Vid. S2**).

Finally, phagocytosis of Aβ and subsequent compaction into vesicles was observed with BAS-exposed microglial cells (**Fig. 1K**). However, when interacting with SH-SY5Y neurons, Aβ adopted a fluffy, cloud-like morphology. Aggregates of Aβ were observed bound with the membrane of SH-SY5Y neurons.

### Clusters of BAS and Aβ are formed inside microglia and released to cause toxicity and depletion on bystander neuronal cells in vitro

We examined clusters composed of BAS and Aβ within phagocytic compartments of microglia and assessed how the convergence of these two species may impact nearby neurons (**Fig. 2A**). BAS and Aβ were tagged with fluorescent fluorophores and sequentially exposed to microglia. Confocal microscopy confirmed colocalization of BAS and Aβ inside microglia (**Fig. 2B**). This colocalization was further validated by high Pearson correlation coefficient values (**Fig. 2C**).

**Fig. 2:**
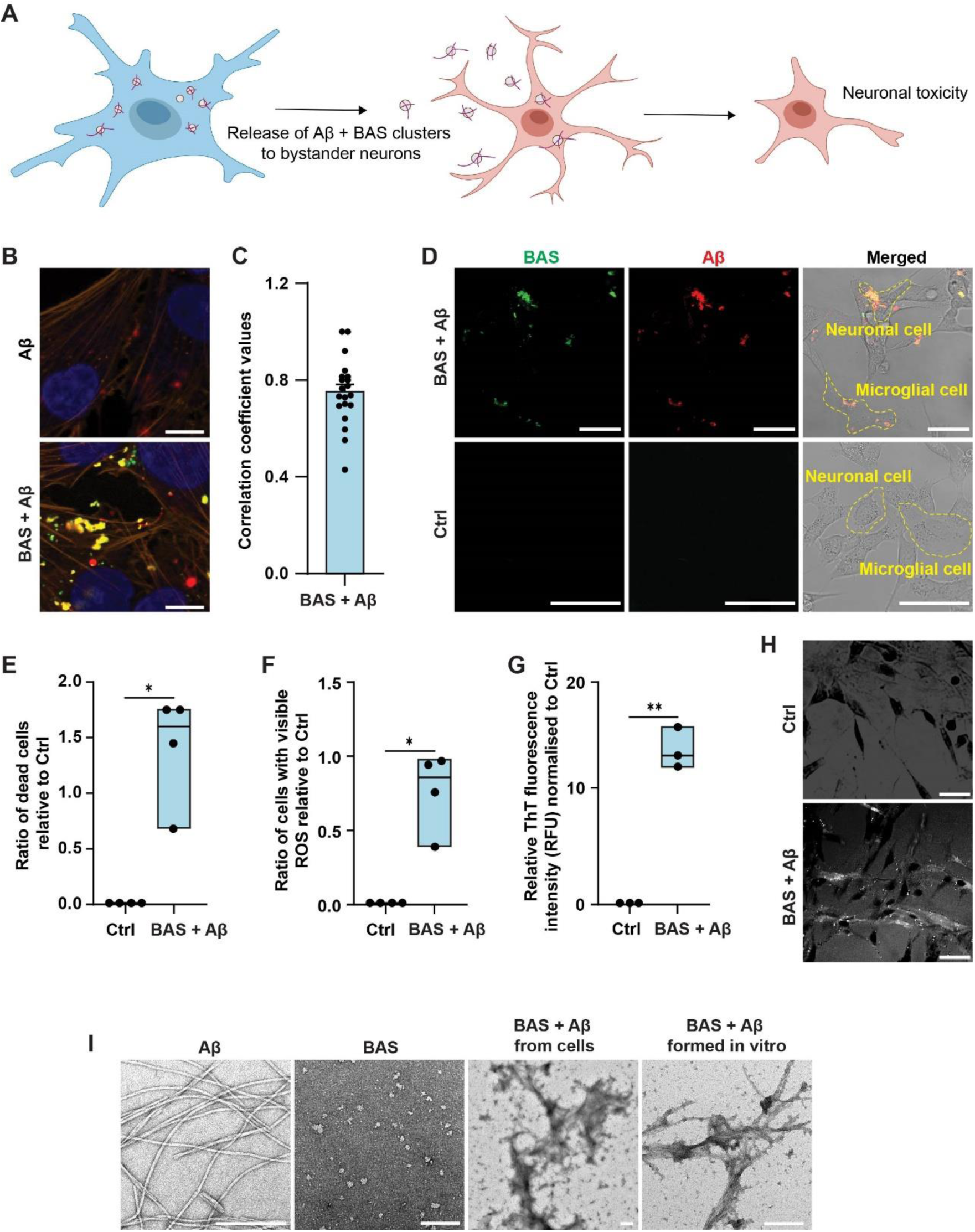
Microglia exposed to BAS followed by Aβ promote neurotoxicity through poorly performed phagocytic degradation leading to regurgitation of clusters that prove toxic to neurons. **A**) Illustration demonstrating microglia regurgitating clusters of BAS and Aβ and incite neurotoxicity. **B**) Representative immunofluorescence image showing colocalization (yellow) of BAS (green) and Aβ (red) inside the phagocytic compartments of HCM3 cells (scale bar 10 µm). **C**) Graph showing Pearson’s correlation coefficient as a quantitative measure of colocalization (GraphPad Prism software analysis) n=20 cells. **D**) Representative immunofluorescence, bright field overlay image of BAS (green) and Aβ (red) clusters (yellow) being regurgitated by microglia and sticking to SH-SY5Y cells in a co-culture environment (scale bar 50 µm). **E**) Graph of measured cytotoxicity for SH-SY5Y using Propidium iodide staining (24 h post exposure) is significantly greater than untreated control (*P ≤ 0.05) following exposure to expelled clusters of BAS + Aβ. **F**) Graph shows ROS activity in SH-SY5Y cells following exposure to BASBAS, Aβ expelled clusters is significantly greater than untreated control (*P ≤ 0.05). **G**) Graph demonstrating the presence of amyloid structures in clusters measured through amyloid fluorescence dye Thioflavin T (ThT). ThT fluorescence was significantly higher in microglia containing dual species clusters relative to untreated control (**P ≤ 0.01). **H**) Polarised light microscopy further confirming β-sheets birefringence of BAS + Aβ clusters from HMC3 cells (scale bar 200 µm). **I**) Transmission electron microscopy (TEM) images comparing the morphology of single BAS or Aβ, as well as clustered species prior to and following expelled. Gross morphological changes observed with clustered species (scale bar 200 nm). All The data are representative of *n* = 3 independent experiments. Statistical values are calculated by one sample t test.

To assess the impact of BAS + Aβ activated microglia on neuronal toxicity, we performed co-culture experiments whereby HMC3 microglial cells exposed to BAS + Aβ were cultured with untreated SH-SY5Y neurons. Time-lapse imaging over 12 hours revealed that fluorescently tagged BAS + Aβ were clustered together and co-expelled by HMC3 microglial cells and actively interacted with SH-SY5Y cells (**Vid. S3**). To further extend our live cell imaging results, confocal microscopy also demonstrated that BAS + Aβs formed co-localising clusters inside HMC3 cells, which subsequently became exocytosed and formed an observable aggregation on the outside of SH-SY5Y neuronal membranes (**Fig. 2D**).

To determine whether the coordinated expulsion of BAS + Aβ was harmful to SH-SY5Y neurons, we performed cytotoxicity and ROS assessments on SH-SY5Y cells co-cultured with BAS + Aβ-exposed HMC3 cells. Results from propidium iodide cytotoxicity assays showed a significant increase in the ratio of dead cells relative to SH-SY5Y cells co-cultured with unchallenged HMC3 microglia (**Fig. 2E**). Additionally, the ratio of cells with detectable ROS was significantly higher in SH-SY5Y neurons co-cultured with BAS + Aβ-challenged HMC3 cells (**Fig. 2F**).

To examine the amyloid structure of regurgitated clusters, we exposed HMC3 microglia to BAS + Aβ and subsequently analysed the cell lysates using Thioflavin T (ThT) fluorescence. Compared to the untreated control, higher ThT intensity was observed in the lysate of BAS + Aβ-exposed cells (**Fig. 2G**). Polarised light microscopy further confirmed the β-sheets birefringence of BAS + Aβ clusters from HMC3 microglial cells (**Fig. 2H**). Transmission electron microscopy (TEM) analysis of BAS + Aβ clusters collected from HMC3 lysates demonstrated fibrillar morphology, which was consistent with that observed when BAS + Aβ where imaged in the absence of cells (**Fig. 2I**).

Finally, we assembled *in vitro* clusters of BAS + Aβ amyloid fragments and directly exposed them to SH-SY5Y cells. We monitored cytotoxicity and ROS levels (**Fig. S3**) and confirmed comparable results to those obtained from HMC3-expelled clusters (**Fig. 2E, F**).

### Proteomics signature of microglia with BAS and Aβ exposure are associated with neurodegeneration and a hyper-active phenotype

To investigate differentially modulated proteins in the context of BAS and Aβ exposure, we collected protein lysates from HMC3 microglia previously exposed to BAS + Aβ, BAS or Aβ, and performed high-throughput proteomics. We specifically focused on proteins in the BAS + Aβ group that showed a significant magnitude of change (Log2 fold change) greater than in the BAS or Aβ-only groups. All examined proteins had adjusted p values greater than 1×10^−5^ (**Fig. 3A**).

**Fig. 3:**
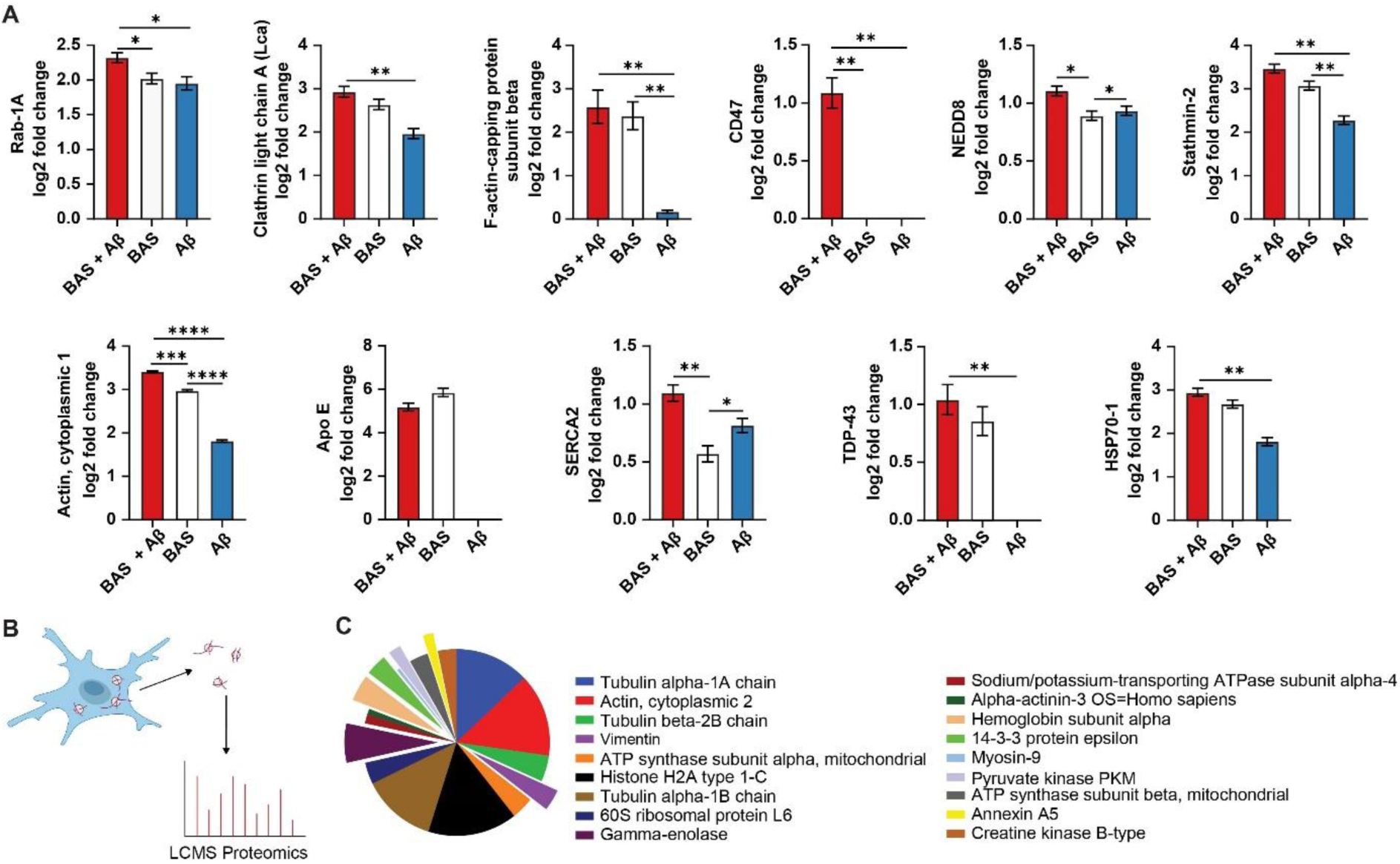
Bacterial amyloid seeds (BAS) and Aβ clustering inside of phagocytic compartments increases expression of proteins linked to neurodegeneration and microglia hyperactivity. **A**) Graphs showing individual proteins significantly increased in cells exposed to BAS and Aβ relative to single species. Eleven proteins linked to AD pathology or microglia hyperactivation were identified as upregulated in HCM3 cells exposed to BAS and Aβ relative to either BAS or Aβ as lone agents (*P ≤ 0.05, ** P ≤ 0.01, *** P ≤ 0.001, **** P ≤ 0.0001). Data are representative of *n* = 3 independent experiments. Statistical values are calculated by unpaired t test. **B**) Illustration demonstrating collection of Aβ + BAS clusters from inside of phagocytic compartments followed by proteomics analysis of proteins adsorbed to the surface of the clusters. **C**) Pie chart illustrating eighteen identified proteins adsorbed to clusters with a critical false discovery rate of 1.0% (ProteinPilot software). Wedges cut from the pie are for proteins previously shown to be related to activated microglia, influence chemokine regulation, or impact AD pathology.

Rab-1A, a known trafficking protein and regulator for autophagy^22^, was significantly upregulated in the BAS + Aβ group. Clathrin light chain A, an endocytic intracellular transport protein involved in cell migration and actin recruitment^23^, was also noticeably higher in the BAS + Aβ group, as were filamentous and cytoplasmic actin - proteins associated with adhesion, shape, division, and migration.^24^ In particular, F-actin, associated with phagocytosis, was greatest in the BAS + Aβ group, relative to the single agents. Apolipoprotein E, an established genetic risk factor for AD^25^, resulted in a greater than five log2 fold increase in the BAS + Aβ, and BAS-only group comparisons. Conversely, no such increase was observed for the Aβ-only group, compared to untreated microglia.

Calcium-ATPase type 2 in the sarco-/endoplasmic reticulum (SERCA2), previously identified in activated microglia in the brains with Alzheimer’s disease patients^26^, was markedly increased in the BAS + Aβ group relative to either the BAS or Aβ-alone counterparts. This was also the case for the pro-inflammatory protein, neuronal precursor cell-expressed developmentally down-regulated protein 8 (NEDD8),^27^ known to participate in the formation of ubiquitinated inclusions and neurofibrillary tangles^28^.

We further found increased expression of Stathmin-2, and TDP-43 in the BAS + Aβ group, both of which are components of insoluble cytoplasmic aggregates.^29^ TDP-43 pathology is an established neuropathological feature in a large number of AD cases.^30^ HSP70-1, which is known to be increased in the presence of oxidative stress and is considered a damage-associated molecular pattern (DAMP)^31^, was significantly overexpressed in the BAS + Aβ group compared to the Aβ-only group, and marginally more so than in the BAS-only group. Finally, the neuronal transmembrane protein CD47, a known “don’t eat me” signal protein, was upregulated in the BAS + Aβ group. Interestingly, none of the other two groups showed upregulation of this important antiphagocytic protein.

### Protein interactome of BAS + Aβ clusters expelled by microglia encourage a hyper-active phenotype and are linked to AD pathology

Given the toxic effect of regurgitated clusters on SH-SY5Y neuronal cells, we performed proteomics to analyse the interactome of the cluster surface (**Fig. 3B**) and assess whether adsorption of specific proteins may be influencing AD pathogenesis. Clusters of BAS + Aβ isolated from HMC3-challenged lysates were purified *via* ultracentrifugation to remove the unbound cellular protein and debris (**Fig. S4**). Eighteen proteins with a false discovery rate of 1.0% were identified using ProteinPilot software (**Fig. 3C**).

While most of the identified proteins were involved in cytoskeletal rearrangements associated with phagocytosis, several proteins stood out given their previously established roles in activated microglia, chemokine regulation, or AD pathology. Notably, the presence of vimentin, a hallmark of reactive astrocytes was of particular interest, as was neuron-specific enolase (Gamma-enolase)^32^, previously identified through a meta-analysis as being elevated in patients with AD.^33^ Haemoglobin subunit alpha and 14-3-3 protein epsilon, both known to interact with Aβ and promote aggregation^34,35^, were also found adsorbed to the expelled clusters of BAS + Aβ. Additionally, pyruvate kinase (PKM) a known promotor of neuroinflammation^36^ was identified. Lastly, Annexin A5, previously recognised as a disease severity marker in the plasma of AD patients^37^, was also found on the surface of the clusters.

### Apoptosis-associated speck-like protein containing a CARD (ASC) speck formation increases following exposure to BAS + Aβ and exacerbates Aβ plaque formation in human monocyte-derived microglia-like cells (MDMi)

NLRP3 is a cytosolic pathogen recognition receptor (PRR) predominantly expressed in microglia in the CNS^38^, it detects a wide range of stimuli and initiates an inflammatory response by forming a multi-protein complex called inflammasomes^39^. Upon stimulation, NLRP3 recruits the adaptor protein ASC and caspase-1. The activation of caspase-1 leads to the release of IL-1β. During NLRP3 activation, ASC forms a perinuclear punctate structure, approximately 1 µm diameter, known as “speck” ^40,41^. Given the importance of inflammasomes in the release of proinflammatory cytokines and following our earlier observation of increased NLRP3 expression (**Fig. 1F**), we sought to determine if NLRP3 inflammasome machinery was activated, as indicated by increased ASC speck formation and IL-1β release upon BAS + Aβ exposure, compared to BAS or Aβ alone.

To this end, we first monitored ASC speck formation using inflammasome reporter cells: THP-1-derived-macrophages (THP1-ASC-GFP) stably expressing an ASC-GFP fusion protein under the control of an NF-κB-binding promoter. In an attempt to fully recapitulate human adult microglia responses, we also generated primary human monocyte-derived microglia cells (MDMi) following an established protocol and staining against ASC to elicit immunofluorescence^42,43^. BAS and Aβ were tagged with AF555 (red) and AF488 (green) fluorescence fluorophores, respectively, prior to ensuing delivery into MDMi or THP1 cells. A schematic of the protocol developed for ASC speck analysis is shown in **Fig. 4A**.

**Fig. 4:**
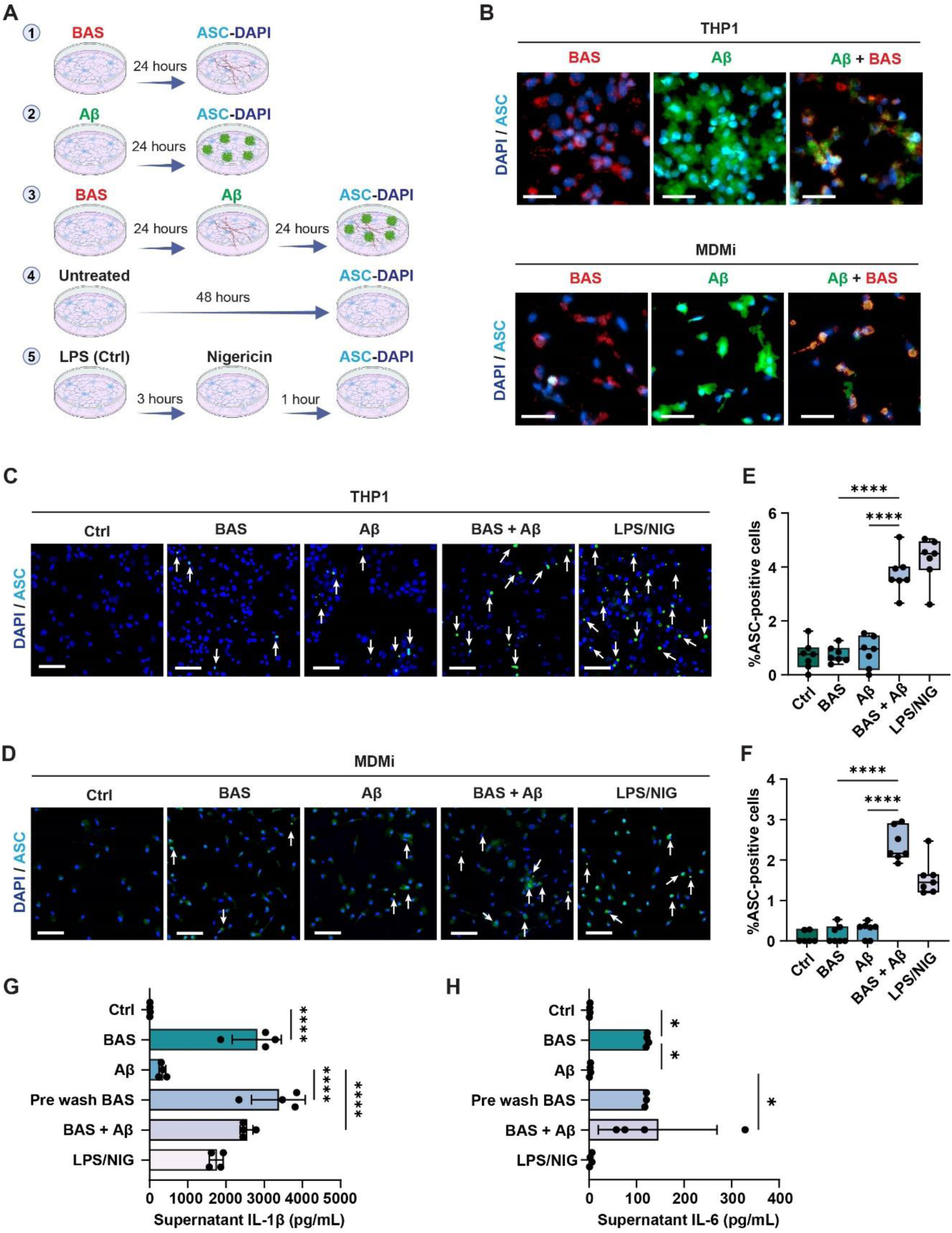
Microglia experience heightened inflammasome activation and pro-inflammatory cytokine release following exposure to BAS and Aβ relative to single species exposure. **A**) Schematic describing experimental workflow in human-microglia-like (MDMi) cells and mature macrophages developed from phorbol 12-myristate-13-acetate treatment of THP-1 monocytes. THP-1 inflammasome reporter cells (THP1-ASC-GFP). **B**) Representative immunofluorescence images confirming colocalization events between BAS (red) and Aβ (green) in THP-1 and MDMi, respectively (scale bar 200 µm). **C, E**) In THP-1 cells, Apoptosis-associated speck-like protein containing a CARD (ASC) speck formation (green) was much higher than in any other groups (**** P ≤ 0.0001). **D, F**) In MDMi cells, speck formation was significantly greater (**** P ≤ 0.0001) in the BAS followed by Aβ group relative to single species exposure (scale bar 20 µm). **G, H**) Graphs demonstrating increased secretion of pro-inflammatory cytokines (IL-1β and IL-6) in the media of MDMi cells fed BAS followed by Aβ relative to single species exposure (**** P ≤ 0.0001, *P ≤ 0.05, respectively). Data are representative of *n* = 3 independent experiments. Statistical values are calculated by One-way analysis of variance (ANOVA).

We found that, Aβ fibrils alone, appeared as fluffy aggregates on the THP1 and MDMi cells. However, in the sample where cells were pre-exposed to BAS, Aβ was internalised and corralled into smaller intracellular aggregates (**Fig. 4B**). Size analysis of Aβ alone versus Aβ + BAS showed greater Aβ fibril compaction in both THP1 and MDMi cells (**Fig. S5**). This observation extends our findings from HMC3 cells, indicating that BAS treatment promotes the phagocytosis of larger Aβ plaques by human microglia (MDMi) and macrophages (THP1).

ASC speck formation was visible in all three samples-Aβ, BAS and Aβ + BAS; however, we identified a synergistic accumulation of ASC specks in the BAS + Aβ group that exceeded the levels of Aβ or BAS alone (**Fig. 4C, D**). Indeed, the number of ASC speck-positive cells were highest in the Aβ + BAS sample in both cell populations (**Fig. 4E, F**). To confirm NLRP3 inflammasome activation in microglia, we examined the presence of pro-inflammatory cytokine IL-1β secreted in the media of the MDMi cells. We found a significant increase of IL-1β and also additional pro-inflammatory cytokine, IL-6 in the BAS + Aβ group relative to BAS or Aβ alone (**Fig. 4G, H**).

Collectively, our findings support the concept that BAS + Aβ challenge in adult human microglia and macrophages result in strong NLRP3 inflammasome activation, leading to ASC speck formation and pro-inflammatory cytokine release.

### Human brain organoids injected with BAS +Aβ show colocalization between the two species, promoting microglial activation and migration towards Aβ

Induced pluripotent stem cell-derived brain organoids have become an important platform for modelling the human brain, as they accurately recapitulate complex neuronal networks.^44^ To extend our previous experiments into a more physiologically complex microenvironment, we utilised a brain organoid model containing functional microglia.^45^ Organoids were microinjected with BAS + Aβ and incubated for one week (**Fig. 5A**). Immunoreactivity was shown through colocalization of Aβ with high expression of protein, Immune-associated gene 1 (Iba1) – characteristic of activated microglial cells, with a Pearson correlation coefficient 0.75 (**Fig. 5B, C**).

**Fig. 5:**
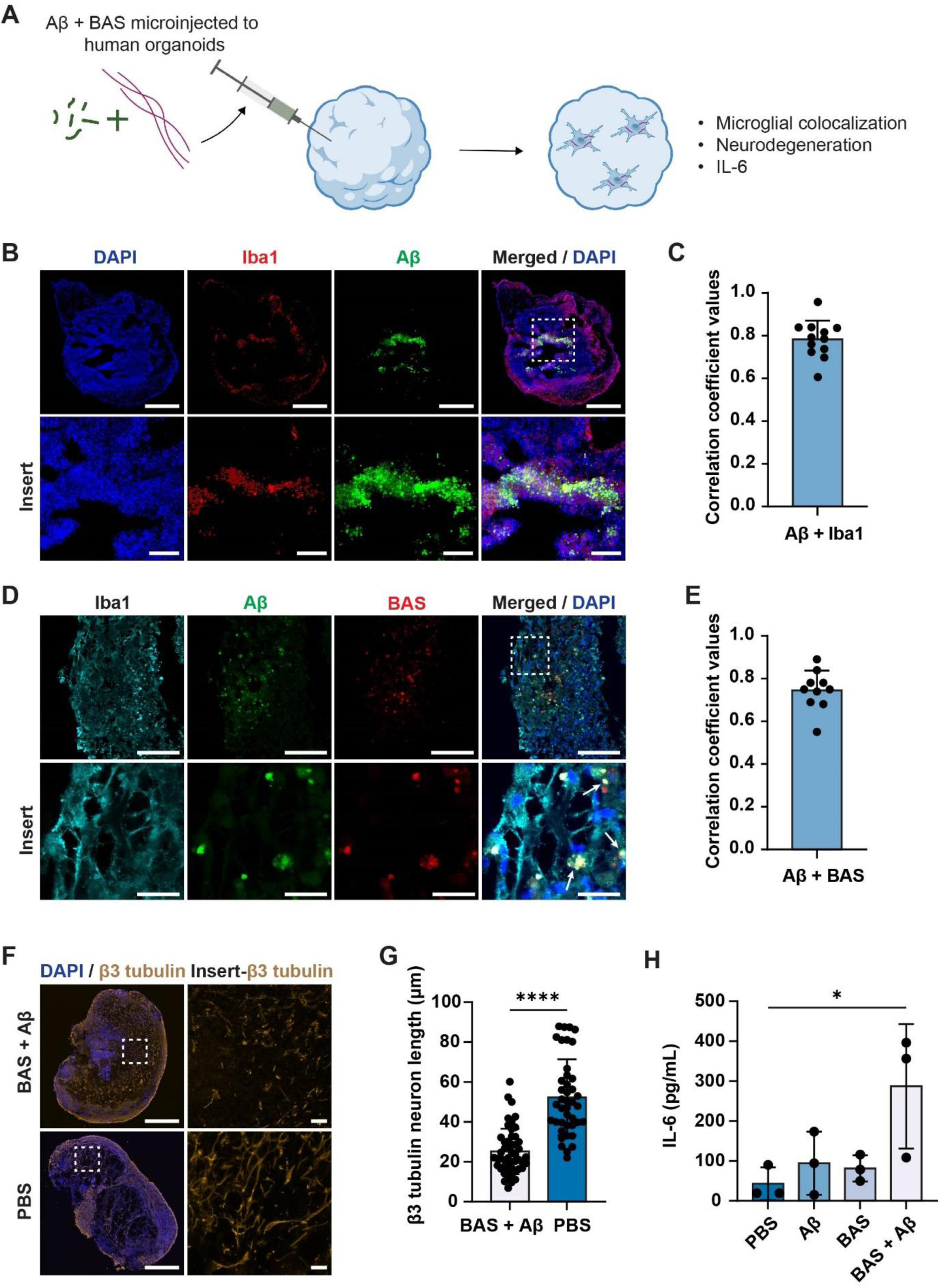
Microbial amyloids seeds (BAS) and Aβ clusters are identified within activated microglia and associated with neuronal shortening and pro-inflammation in an immunocompetent human brain organoid model. **A**) Schematic visualisation of stem cell derived brain organoids injected with BAS and Aβ prior to performing analysis of colocalization, neurodegenerative phenotype and pro-inflammatory cytokine secretion. **B**) Representative immunofluorescence image of brain organoid marking quantifiable colocalization between microglia measured by Iba1 (red) and Aβ (green) (scale bar 400 µm; insert: 100 µm). **C**) Graph of correlation determined by Pearson’s correlation coefficient (GraphPad Prism software). **D**) Representative immunofluorescence image demonstrating a triad of localisation occurring between microglia (cyan), Aβ (green) and BAS (red) (scale bar 100 µm; insert: 20 µm). **E**) Graphed confirmation of BAS and Aβ convergent clusters determined by Pearson’s correlation coefficient (GraphPad Prism software). **F**) Representative immunofluorescence image showing the pattern of beta (β)3 tubulin protein expression (gold) - as a marker for neuronal health is altered in brain organoids exposed to BAS and Aβ relative to phosphate buffered saline treated control (PBS) (scale bar 500 µm; insert: 25 µm). **G**) Graphed data demonstrating a decrease in β3 tubulin length as a measure of neuronal health following exposure to BAS and Aβ (quantified immunofluorescence data) relative to PBS (**** P ≤ 0.0001). **H**) Graphed data for IL-6 expression in the organoids injected with BAS + Aβ, Aβ and BAS. IL-6 expression was significantly (*P ≤ 0.05) higher in organoids injected with BAS + Aβ in comparison to PBS control, Aβ or BAS alone. All The data are representative of *n* = 3 independent experiments. Statistical values are calculated by One-way analysis of variance (ANOVA).

To further assess the coordinated clustering and colocalization of BAS and Aβ within microglia phagosomes, we injected the organoids with fluorescently labelled BAS (red) and Aβ (green). We then examined microglia positioning and morphology in relation to BAS and Aβ and observed similar morphological patterns compared to our 2D *in vitro* cultures. This suggest that Iba1-positive (activated) microglia (cyan) contained colocalised BAS and Aβ clusters (**Fig. 5D, E**) with a Pearson correlation coefficient 0.78).^46^

We also analysed the neurodegenerative effects of BAS + Aβ clusters on organoids neurons using β3 tubulin immunoreactivity as a proxy for neuronal health (**Fig. 5F**). The average length of β3 tubulin-positive neurons was reduced by over 50% from 52.32 (PBS) to 25.16 µm in BAS + Aβ treated organoids (**Fig. 5G**). Additionally, IL-6 levels increased more than six-fold, from 42.96 pg/mL (PBS) to 287.12 pg/mL in the BAS + Aβ-treated organoids (**Fig. 5H**).

These organoid experiments support our cellular model findings, suggesting that the convergence of BAS and Aβ within the phagocytic compartments of microglia leads to microglial activation and the propagation of a pro-inflammatory phenotype.

### Zebrafish orally fed with BAS and cerebrally microinjected with Aβ exhibited translocation of both species to the brain, resulting in impaired synaptic function and cognitive decline

To investigate the interplay between BAS and Aβ in the central nervous system (CNS) and the resulting neurotoxic events in an *in vivo* setting, we utilized adult zebrafish. Zebrafish were orally fed with BAS and/or cerebrally microinjected with Aβ. We then examined histopathological changes and cognitive decline outcomes (**Fig. 6A**).

**Fig. 6:**
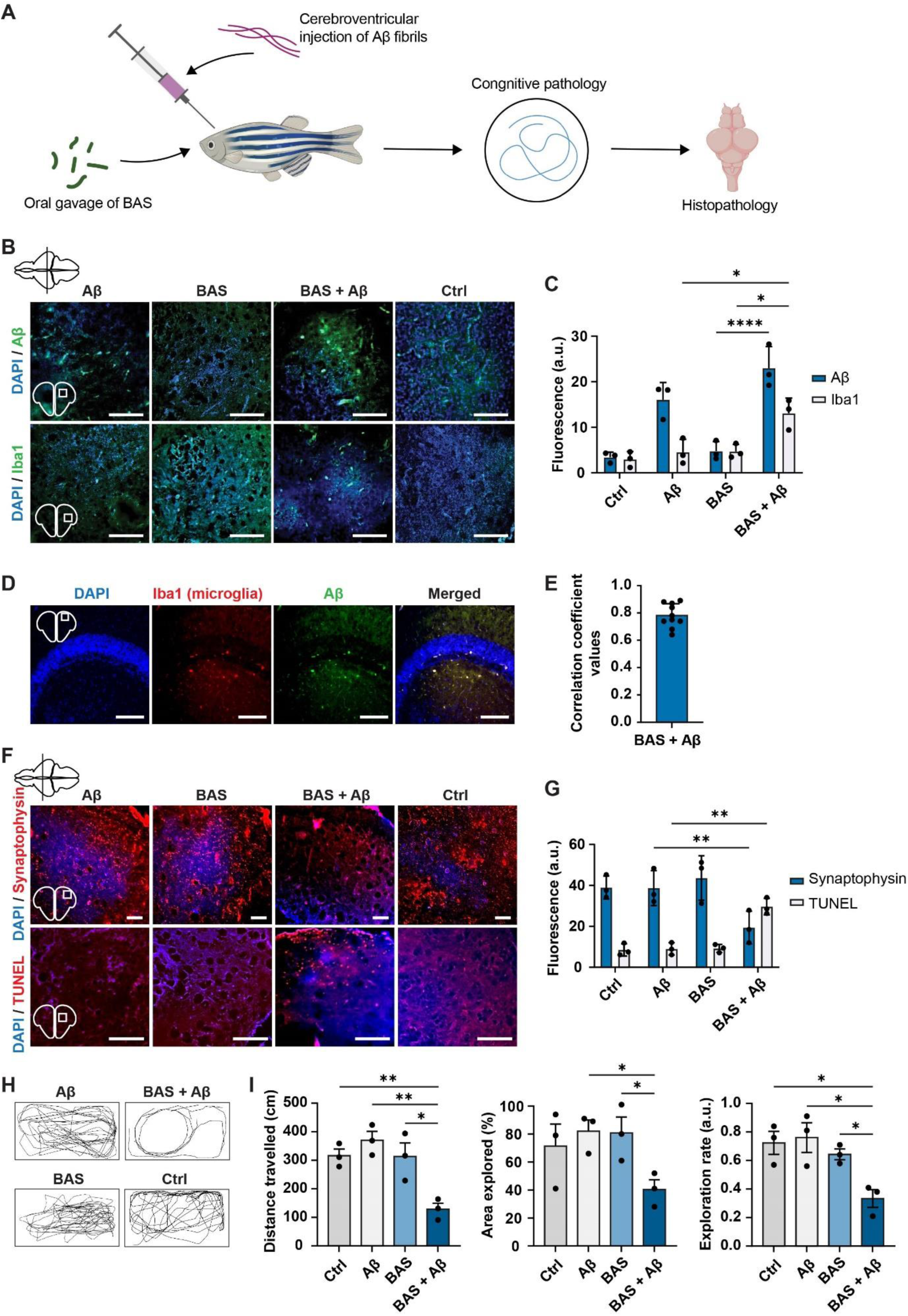
Orally fed microbial amyloids and cerebroventricular injected Aβ converge in the brain of zebrafish larvae leading to marked cognitive decline. **A**) Schematic overview of experimental procedure to microinject adult zebrafish brain with Aβ and orally administer BAS. **B**) Representative immunofluorescence image of zebrafish brain, demonstrating increased Aβ expression (green) in fish exposed to BAS + Aβ relative to single Aβ or BAS alone (scale bar 100 µm). **C**) Quantitation of fluorescence intensity as a marker for Aβ abundance in brain sections. Fluorescence signal is increased when fish is administered with Aβ + BAS relative to Aβ or BAS alone (* P ≤ 0.05, **** P ≤ 0.0001, two-way ANOVA). **D**) Representative immunofluorescence images brain sections of zebrafish demonstrate microglia colocalization with Aβ within this area associated with Alzheimer disease pathology (scale bar 100 µm). **E**) Graphed correlation determined by Pearson’s correlation coefficient (GraphPad Prism software). **F**) Representative immunofluorescence images illustrating changes to synaptophysin and expression and presence of DNA fragmentation (TUNEL) as a marker of neuronal toxicity following exposure to BAS + Aβ relative to single species exposure (scale bar 50 µm). **G**) Quantification of fluorescence intensity used as a marker for the health of neuronal cells, show that synaptophysin expression is significantly decreased, while DNA fragmentation is significantly increased (** P ≤ 0.01, two-way ANOVA). **H**) Movement trajectories of zebrafish following exposure to BAS + Aβ relative to single species. Fish movement is greatly repressed in the BAS + Aβ exposure group relative to other groups. **I**) Graphs representing total distance travelled between groups (5 min observation time). Distance is significantly reduced in BAS + Aβ relative to single species exposure (* P ≤ 0.05, ** P ≤ 0.01). Graphed data referencing percentage area explored shows ∼50% reduction in area reduced in the BAS + Aβ exposure group relative to other groups (* P ≤ 0.05). Graphed exploration rate reduced significantly compared to other groups (* P ≤ 0.05). For behavioural analysis, 6 fish per group and three measurements per fish were recorded in three different times of a day. Statistical values are calculated by unpaired t test.

First, we performed immunofluorescence on brain slices from zebrafish in each group (Aβ, BAS, BAS + Aβ and control) to assess Aβ aggregation and identify microglia co-immunolabeling and reactivity, as indicated by increased Iba1 expression. In the Aβ-only group (green), Aβ was visible as small plaques. However, in the presence of BAS, larger, globular Aβ plaques were observed, accompanied by higher Aβ immunoreactivity (**Fig. 6B, C**). An increased Iba1 signal indicated greater microglial activation in the BAS + Aβ group. Notably, Iba1-positive microglia were colocalised with Aβ (Pearson correlation coefficient 0.781) and showed higher immunoreactivity compared to the Aβ-only group (**Fig. 6D, E**).

Next, we examined the potential neurodegenerative effect of BAS + Aβ clusters by assessing synaptophysin degeneration and *bona fide* DNA damage. Zebrafish that received BAS orally and cerebral microinjection of Aβ displayed the most significant loss of synaptophysin expression compared to Aβ, BAS and mock-treated control groups (**Fig. 6F, G**). Similarly, TUNEL assay revealed a higher incidence of DNA double-strand breaks in the fish that received both BAS and Aβ.

We also monitored cognitive performance by studying swimming behaviour over a 5-minute period. Notably, the zebrafish that received BAS orally and Aβ via cerebral microinjection exhibited the greatest decline in swimming trajectory behaviour (**Fig. 6H**). We quantified overall behaviour in terms of distance travelled, percentage of area explored, and exploration rate. While the BAS- and Aβ-only groups showed a similar pattern of behaviour to the buffer control group in each of the three experiments, the BAS *+* Aβ group demonstrated a significant decline in all categories. Instead of the erratic, fast paced movement observed in the other groups, zebrafish in the BAS *+* Aβ group had a propensity to move in a slow circular pattern, which significantly decreased the distance, area, and exploration rate. The amount of distance travelled overtime was reduced almost 4-fold the in BAS + Aβ sample. Similarly, the area explored, and exploration rate was reduced by 53% in the BAS + Aβ-treated animals (**Fig. 6I**).

## Discussion

Alzheimer’s disease (AD) is a multifactorial and progressive neurodegenerative disorder. While Aβ aggregation has traditionally been considered central to its pathophysiology, other mechanisms such as ApoE trafficking, the formation of intraneuronal Tau tangles, mitochondrial dysfunction, and microglial-driven neuroinflammation are becoming increasingly recognised as contributors to disease progression.^47^ Despite these insights, most therapeutic strategies to date have focused on these pathological markers individually, resulting in limited success in improving cognitive function in clinical trials.^48^

Recent evidence highlights the potential role of gut dysbiosis and the gut-brain axis in influencing AD onset and severity.^2,49^ However, the putative molecular triggers derived from gut pathogens and their mechanism of entry into the brain, where they might modulate AD progression, remain poorly understood. We and others have previously shown that microbial amyloids, such as FapC from *P. aeruginosa* and and CsgA from *E. coli,* can directly accelerate the aggregation Aβ and alpha-synuclein (αS) and exacerbate their neurotoxic effect.^6,7^ These findings suggest that microbial-derived amyloids could be a key link between guts dysbiosis and AD pathogenesis, opening a new avenues for understanding and potentially intervening in the disease process.

Based on the cross-β sheet architecture of amyloids, it is reasonable to hypothesize that microbial amyloids can access the brain in a prion-like mechanism as previously demonstrated with αS amyloids.^50^ Moreover, microbial amyloids may access the brain through extraintestinal infections, including those affecting the nervous system. Under these circumstances, microbial biofilms, along with embedded amyloids, could feasibly interact with immune microglia. This is especially relevant given the observation of pro-inflammatory microglia around Aβ plaques in AD brains, and the modulation of microglial subpopulations by the gut microbiome,^51,52^

It is therefore hypothesised that microbial amyloids originating from the gut could exacerbate AD pathology by interacting with Aβ, triggering microglial hyperactivity-dependent inflammation. Gut microbes are known to form interspecies biofilms, using their amyloids as a cross-seeding agents.^20^ To model gut dysbiosis and the potential pathogenic effects of interspecies biofilms, we used a combination of FapC and CsgA amyloids (BAS) from *P. aeruginosa* and *E. Coli* for this study. Using a variety of biological models, we tested the hypothesis that a toxic, synergistic interaction between BAS and Aβ exists, which drives a reactive microglial phenotype, leading to progressive neurotoxicity.

Our findings demonstrate that, compared to BAS or Aβ alone, the combination of BAS + Aβ clusters rapidly polarized microglia to a pro-inflammatory state. Proteomic profile of microglia exposed to BAS + Aβ identified the upregulation of clathrin light chains, actin cytoplasmic, and Stathmin-2. These proteins are associated with neurodegeneration and have been detected in early stages of AD pathology.^29,53^ We further identified a marked increase in ApoE expression in the BAS + Aβ group. ApoE is a well-known genetic risk factor for AD ^54^, and its implicated in promoting a neurodegenerative microglial phenotype.^55,56^

Additionally, our data suggests that the combination of BAS and Aβ clusters enhances AD pathology by increasing expression of the NEDD8, a protein linked to inflammation and can potentially encourage the formation of ubiquitinated inclusions in the brain ^27,28^. Further, we propose that clusters of BAS + Aβ-induced reactive microglia, much like cancer cells, may express elevated levels of CD47, which acts as a way of overriding healthy mechanisms of apoptosis by astrocytes in the presence of aberrantly behaving cell populations ^57^.

Next, we established those toxic cellular interactions between microglia and neurons occurred as a result of BAS + Aβ clusters. We observed that the initial BAS + Aβ clusters formed within microglia were later regurgitated, retaining a distinct interactome, which was visible under TEM. The importance of this finding is best echoed in human study data that shows small Aβ aggregates are most toxic and immunoinflammatory, often appearing in early-stage AD brain across various regions.^58^

Among the proteins we found adsorbed to the regurgitated clusters, some were pro-inflammatory while others were directly linked to AD pathology.^34,35,37^ For instance, vimentin, a marker of inflammation was present on these clusters. Vimentin, together with activated astrocytes, is known to be associated with cellular hypertrophy and the excessive release of pro-inflammatory cytokines.^59^ This suggest that vimentin could contribute the inflammatory responses seen in AD brains.

Another significant finding was the presence of pyruvate kinase (PMK), a protein linked to metabolic changes in AD and enhancer of gamma-secretase activity, which plays a role in the production of Aβ. ^60^ PMK, specifically its isoform PKM2, has also been implicated in inflammatory processes through its promotion of complement component 1q (C1q), tumor necrosis factor-alpha (TNF-α), and interleukin-1 alpha (IL-1) secretion.^61^ The knockdown of PKM2 in previous studies was shown to inhibit microglia activation and reduce the inflammatory response, suggesting that PKM plays a pivotal role in maintaining neuroinflammatory states in AD.^61^

ASC specks formation can be used as a simple upstream readout for inflammasome activation.^40,62^ Commonly detected in the extracellular milieu, ASC specks are a regular occurrence in a range of inflammatory diseases and are generally associated with poor prognosis.^63^ In AD, elevated levels of ASC speck have been observed in the cerebral spinal fluid (CSF) where they can exacerbate pathology by cross-seeding the aggregation of Aβ.^41,64^

To explore this in more detail, we used human MDMi and THP1-ASC-GFP cells to monitor ASC speck formation and its colocalization with Aβ, BAS and BAS + Aβ. Our results showed that sequential exposure to BAS + Aβ resulted in the highest level of ASC speck formation relative to BAS, or Aβ alone, indicating a synergistic effect. This was further supported by a significant increase in the secreted inflammasome pro-inflammatory cytokine IL-1β in cells exposed to BAS + Aβ compared to either Aβ or BAS alone.

These findings align with previous research by Venegas et al. who demonstrated that ASC specks generated by microglia can bind to Aβ, promoting the formation of Aβ oligomers and aggregates which are crucial in AD progression^65^. Previously, we have observed that FapC can accelerate the fibrilization of Aβ and exaggerate its toxicity *in-vitro* and *in-vivo* in zebrafish models.^7^ CsgA and culri-expressing *E. coli* have shown to increase the α-Synuclein pathology both in the gut and brain of mice models.^6,8^ Aβ also disintegrated the architecture of *P. aeruginosa* and *E. coli* biofilms by remodelling FapC and CsgA amyloids, indicating a role of amyloid-proteins in bidirectional communication across gut-brain axis.^66^ These studies provided an insight into biophysical interaction and direct cross-seeding between microbial amyloids and neuronal amyloid proteins Aβ and α-Synuclein. Taken together, our results provide an understanding of the pathological interplay between BAS, Aβ and microglial hyperactivation. This “pathological trifecta” highlights a mechanism by which BAS and Aβ work in concert to hyper stimulate microglia, potentially accelerating neuroinflammation and contributing to the poor prognosis commonly observed in AD.

In brain organoids, we further observed clear clustering of BAS with Aβ, accompanied by an activated microglia phenotype. This is consistent with previous studies that link inflammation in the brain to neurodegenerative processes.^67–69^ The neurodegenerative effect of the combined BAS + Aβ challenge were evident in the form of fragmented and shortened neurons, pointing to a significant detrimental impact on neuronal integrity. This aligns with observations in AD, where neuronal damage and inflammation are key pathological features.

To further assess the *in vivo* relevance of this findings, we employed a zebrafish model. This allowed us to trace each toxic event from the colocalization and clustering of BAS and Aβ to subsequent microglial activation, neuronal cell death, and cognitive decline, ultimately showcasing a neurodegenerative phenotype. Using synaptophysin as a marker for synaptic integrity, we found a significant reduction in synaptophysin levels in zebrafish exposed to BAS + Aβ mirroring findings in AD brain tissue, where synaptic loss is a hallmark of disease progression.^70^

Moreover, zebrafish exposed to BAS + Aβ exhibited significantly increased cell death, particularly in neurons, and marked cognitive impairment. These results underscore the potential role of microglia in mediating the toxic effects of BAS + Aβ clusters. The findings suggest that once microglia encounter these clusters, they may drive the neuroinflammatory response, contributing to the progressive onset and worsening of AD.

This study brings a new line of evidence that positions microglia-based progressive inflammation as a potential AD pathway driven by gut microbiome dysbiosis, microbial amyloids, and age-related gut permeability. While it outlines a novel route of AD pathogenesis, future research is critical to delve deeper into the molecular mechanisms that allow pathogenic metabolites from the gut to reach the brain. Key questions remain, such as the specific signals that facilitate the migration of microbial byproducts across the blood brain barrier and how this signals interact with Aβ and microglia once inside the brain.

It may be particularly interesting to look more in depth at the protein interactome known to build up on Aβ *in vivo* and investigate whether the interactome may actually be the driving force that initially promotes BAS to colocalise with Aβ. Moreover, whether this event happens in the gut or following translocation to the brain. Our study is the first to explore the detrimental combination of BAS, Aβ, and hyper-active microglia as a pathological mediator in AD, creating the opportunity to develop intervention strategies as our understanding continues to develop.

## Methods

### Animal Husbandry and Ethics Statement

Experiments with adult zebrafish (wild type AB *Danio rerio*) were performed with 10 months old fish. The fish was maintained before and during the experiment with 28 °C water circulatory system with 14 hrs light and 10 hrs dark cycle at the University of Queensland Seddon Aquarium Biological Resources Facility. The fish was anesthetised with 0.01% tricaine in fish water for 20 sec before microinjection. At terminal tissue collection, head of the fish was excised from trunk. The fish was placed in an ice-chilled 0.4% tricaine (Holtfreter’s buffer) and the head was excised from the trunk with a surgical blade. The excision was performed under a stereomicroscope and the heads were washed thrice in phosphate-buffered saline (PBS, pH 7.4) and then stored in 2.5% paraformaldehyde. All in vivo experiments with zebrafish were performed according to the University of Queensland ethical guidelines and the protocols were approved by the Animal Ethics Committee AEC (2020/AE000382). All other experiments, including in vitro, were performed according to the Occupational Health & Safety (OHS) guidelines of the University of Queensland. Ethical approval for collecting and utilising human donor blood for generating MDMi cells was obtained from The University of Queensland Human Research Ethics Committee (HREC approval #2020000559).

### Microbial amyloid seeds preparation

Bacterial amyloids FapC and CsgA were formed by incubating 1mg/mL concentration of these proteins in PBS at 37 °C for 2 weeks and kept at 4 °C for future experiment. Aβ fibrils were prepared in 0.1% Dimethylsulfoxide (DMSO)/MQ water and incubated at 37 °C for one week. Any potential contamination from endotoxins in FapC and CsgA was removed by Pierce high capacity endotoxic removal spin columns by three washings as per manufacturer instructions. FapC and Aβ were monomerised by incubation hexafluoro-2-propanol (HFIP) for 2 hrs and then drying. CsgA was monomerised by incubation with HFIP/formic acid (1:1) for 2 hrs and dried on the bench. The mixture of the seeds of FapC and CsgA was prepared by mixing FapC and CsgA amyloids (1:1) and probe sonicating the mixture in an ice water bath for 2 minutes on 1 second on/off cycle right before the experiment (Sonics Vibracell: 750 watts and 20kHz, 20% amplitude). The mixture of the seeds was prepared fresh every time before the experimentation.

### Cell culture

HMC3 (CRL-3304), SH-SY5Y (CRL-2266) cells were purchased from ATCC, US. Human monocytic inflammasome ASC speck reporter cells, THP1-ASC-GFP, was purchase in Invivogen (thp-ascgfp) HMC3 were culture in DMEM supplemented with 10% foetal bovine serum (FBS) and SH-SY5Y in DMEM/F12 medium with 10% FBS and 1% Peniciliin/Streptomycin. THP1-ASC-GFP cells were cultures in RPMI-1640 (Gibco, #22400-089) supplemented with 10% heat-inactivated fetal bovine serum (FBS; Gibco, #10082147), 0.1% Zoecin (InvivoGen, #ant-zn-05), and 100 U/ml penicillin-streptomycin (ThermoFisher, #15070-063). Cells were maintained at a density of 3 × 10⁵ cells/ml and passaged upon reaching 1 × 10⁶ cells/ml. Before treatment, cells were differentiated into macrophages by incubating with 0.5 μM phorbol 12-myristate 13-acetate (PMA) (Sigma, #P8139) for 3 hours, followed by 16 h incubation in serum-free medium To generate human monocyte-derived microglia-like (MDMi) cells, monocytes were plated at 1 × 10⁵ cells per well in a 96-well plate under serum-free conditions in RPMI-1640 GlutaMAX (Gibco, #61870036) with 100 U/ml penicillin-streptomycin (ThermoFisher, #15070-063), 2.5 μg/ml Fungizone (Life Technologies), and recombinant human cytokines: M-CSF (10 ng/ml), GM-CSF (10 ng/ml), NGF-β (10 ng/ml), MCP-1 (CCL2, 100 ng/ml), and IL-34 (100 ng/ml; all from Peprotech) as previously described.^42^ Cells were cultured under humidified conditions (37°C, 5% CO₂) for up to 14 days to induce microglial differentiation. All cell lines were maintained in an incubator (37 °C / 5% CO_2_) and routinely screened for mycoplasma infection.

### Human monocyte-derived microglia like cells (MDMi) generation

Peripheral blood mononuclear cells (PBMCs) were isolated from buffy coat blood samples obtained from healthy donors through the Australian Red Cross Blood Service. The buffy coat was diluted 1:1 with phosphate-buffered saline (PBS; Lonza, #181828) and processed using SepMate 50 tubes (STEMCELL Technologies, BC, Canada) following the manufacturer’s instructions. PBMCs were then separated using Lymphoprep density centrifugation (STEMCELL) and collected for further use. Monocytes were subsequently isolated from the PBMCs via positive selection with anti-CD14+ magnetic beads (Miltenyi Biotec) and prepared for differentiation. To generate human monocyte-derived microglia-like (MDMi) cells, monocytes were plated at 1 × 10⁵ cells per well in a 96-well plate under serum-free conditions in RPMI-1640 GlutaMAX (Gibco, #61870036) with 100 U/mL penicillin-streptomycin (ThermoFisher, #15070-063), 2.5 μg/mL Fungizone (Life Technologies), and recombinant human cytokines: M-CSF (10 ng/mL), GM-CSF (10 ng/mL), NGF-β (10 ng/mL), MCP-1 (CCL2, 100 ng/mL), and IL-34 (100 ng/mL; all from Peprotech) as previously described.^42^

### Single HMC3 experiments

HMC3 cells (5 x 10^4^) were seeded in a well of a 6-well plate for 24 hrs and subsequently incubated with 0.1 µM of BAS labelled with Alexafluor (AF) 647 (0.1 µM) in cell media for 24 hrs. Labelling with AF was performed by incubating the protein with the dye for 90 min at room temperature followed by washing in PBS using 10 kDa centrifugal filter. After 24 hrs, the BAS media was removed, cells were washed with PBS (5 min) and Aβ fibrils (1 µM) were added to the same well and the cells incubated for a further 24 hrs. Cells were imaged on the Olympus CKX53 (Japan) using Olympus cellSens Imaging Software.

### Co-culture experiments

HMC3 cells (5 x 10^4^) were seeded in a well of a 6-well plate for 24 hrs and subsequently incubated with 0.1 µM of BAS labelled with AF647, in cell media for 24 hrs. Aβ fibrils (1 µM, labelled with AF488) were added to the same well and the cells incubated for a further 24 hrs. The treated cells were trypsinised and centrifuged for 5 min at 1500 rpm. The cell pellet was resuspended in RPMI medium and the cell suspension added into a well of a 6-well plate seeded with (5 x 10^4^) SH-SY5Y cells the day prior. After 6 hrs of incubation, co-cultured HMC3/SH-SY5Y cells were imaged for 12 h using the PerkinElmer Operetta instrument and results were analysed by Harmony High-Content Imaging and Analysis software.

### Western blotting

HMC3 cells were lysed in radioimmunoprecipitation assay (RIPPA) buffer and protein levels were quantified using a Pierce BCA Protein Assay Kit (Thermo Fisher Scientific). Samples were heated at 95 °C for 5 min in 6x loading buffer before loading into a gel. Membranes were probed with three antibodies: NLPR3 ((D4D8T) from Cell Signaling Technology and LAMP1 (D2D11) XP at a 1:1000 dilution - 5% w/v BSA, 1X TBS, 0.1% Tween 20, incubated overnight at 4 °C. Anti-beta Actin antibody (ab8227, Abcam) at a 1:50,000 dilution 5% w/v Milk, 1X TBS, 0.1% Tween 20, incubated for 1 hr at room temperature. Anti-rabbit and anti-mouse secondary antibodies were used at a 1:10,000 dilution 5% w/v Milk, 1X TBS, 0.1% Tween 20, incubated for 1 h at room temperature (Agilent Technologies).

### ELISA analysis of proinflammatory cytokines

The human IL-6 ELISA kit used to measure IL-6 expression was purchased from Abcam (ab178013). The kit was used as per manufacturer’s instructions. Briefly, 50 µL of media was added to a well of the supplied 96-well microplate, followed by 50 µL of antibody cocktail and the plate incubated on a shaker for 1 hr at room temperature. Following 3x subsequent washes with buffer PT, 100 µL of TMB Development Solution was added to the well and allowed to incubate for 10 min in the dark on a shaker set to 400 rpm. Finally, 100 µL of Stop Solution was added to the well for 1 min and mixed on the plate reader before measuring the optical density at 450 nm using an EnSight plate reader (PerkinElmer).

### Lipid droplet formation

HCM3 cells were grown in a six well plate and administered I00 µM of oleic acid in DMEM I0% FBS for 24h. Cells were fixed (4% PFA) for 20 mins and washed x3 in PBS and 60% isopropanol. Lipid droplets were stained with filtered Oil Red O solution (0.5% in isopropanol) for 20 min. Imaging was performed on an Olympus CKX53 using Olympus CellSens Imaging Software.

### Cytotoxicity and reactive oxygen species (ROS) assays

HMC3 cells were cultured in 24-well plate for 70-80% confluency and incubated with BAS (0.1 µM), full length microbial amyloids (0.1 µM), LPS (100 ng/mL) or 0.01% TritonX (positive control) for 24 hrs. Cytotoxicity assay was performed by adding propidium iodide PI (2 µM) into the cells and incubating for 30 min. Cells were imaged for dead PI staining with PerkinElmer Operetta and live-dead ratio was quantified by Harmony Software. ROS expression was measured in live cells using CellROX™ (Thermo Fisher Scientific). CellROX was prepared as per manufacturer’s instructions. Briefly, following treatment, HMC3 cells were incubated in 5 µM CellROX in cell culture media for 30 min. Live cell confocal microscopy (Leica SP8) was used for image acquisition examining ROS expression 1 hr and 24 hrs post treatment. In the co-culture experiment, HMC3 cells were exposed to 0.1 µM of BAS for 24 hrs, followed by washing with PBS to remove the unbound BAS and additional incubation with 1 µM Aβ fibrils. BAS and Aβ fibrils were labelled with AF as described above. Cytotoxicity and ROS was measured by PI (2 µM) and dichlorodihydrofluorescein diacetate (DCFDA) (2 µM) by analysing the fluorescence in live cells with PerkinElmer Operetta as described above.

### Cell imaging for the interaction of Aβ and BAS

HMC3, SH-SY5Y and co-culture of HMC3 and SH-SY5Y cells were treated with AF labelled BAS or BAS + Aβ as discussed in cytotoxicity assays. The cells were fixed with 4% paraformaldehyde in DMEM/F12 for 30 mins, washed with DPBS and stained with Hoechst and Actin ReadyProbes™ (1µL/mL; 30 min). The fixed cells were imaged with Leica SP8 confocal microscopy. For live cell imaging, co-culture of HMC3 and SH-SY5Y cells was exposed to AF labelled BAS (0.1 µM) for 24 hrs, washed with DPBS and then exposed to AF labelled Aβ (1 µM) for 6 hrs and then placed in PerkinElmer Operetta for automated imaging at predefined time intervals of 10 mins. The images were analysed by Harmony Software and stitched in sequence to study the propagation of BAS + Aβ clusters from HMC3 to SH-SY5Y cells. The size and intensity of the fluorescent plaques and colocalization was quantified by Image J.

### Transmission electron microscopy (TEM)

Aβ fibrils were prepared by incubating aqueous solution of Aβ (Anaspec) at 37 °C for one week. Aβ fibrils, FapC and CsgA were subjected to probe sonication as described above, mixed in a ratio of 0.5:0.5:10 (FapC:CsgA:Aβ) and vortexed at 37 °C for 3 hrs. The TEM imaging was performed by negative staining with uranyl acetate. A dope of the sample was placed on a glow discharged carbon-coated copper grid for 1 min and blotted. The grid was washed with a drop of water twice by blotting and then negatively stained with 1% uranyl acetate for 25 sec. The grid was dried and imaged with a Hitachi HT7700 TEM operating at 120 kV.

### Polarised light microscopy

The HMC3 cells and SH-SY5Y cells were cocultured in an 8-well chamber slide and treated with BAS (0.1 µM) for 24 hrs, followed by washing and then incubation with Aβ (1 µM) for additional 24 hrs. The cells were fixed with 4% paraformaldehyde in DMEM/F12 medium for 30 mins, washed with DPBS twice (5 min each) and stained with 0.5% congo red stain in DPBS for 20 min, washed thrice with DPBS and sealed with DABCO mounting medium. The cells were imaged for polarised light microscopy birefringence with an Olympus BX61 microscope fitted with a polarisation filter. The images are presented as bright-light pseudo colors for the apple-green birefringence.

### Thioflavin T fluorescence measurement

Co-culture of HMC3 and SH-SY5Y cells were exposed sequentially to BAS and Aβ as discussed in cytotoxicity assay for 24 hrs and the clustered BAS + Aβ was extracted from the cells by removing the media and treating the cells with RIPA lysis buffer (100 µL) for 10 min at ice. The extracted mixture was centrifuged to a pellet (10,000 g for 30 min) and pellet was dissolved in DPBS and adjusted to the protein content of 1mg/mL by Pierce BCA protein quantification assay kit as per manufacturer directions. The extracted protein mixture was incubated with ThT dye (0.1 µg/mL) for 2 hrs at 37 °C. The ThT fluorescence reading of the mixture was read at excitation 445 nm and emission 485 nm. Untreated coculture of the cells, treated with same extraction procedure and ThT staining was used as control.

### Proteomics

Microglial cells (5 x 10^4^) were incubated with 0.1 µM of BAS in serum free cell media for 24 hrs. After the designated incubation time, cells were washed and 1 µM of Aβ fibrils were added to the same well and incubated for 24 h. Most of the cell medium were removed from each well and cells were treated with 100 µL of RIPA lysis and Extraction buffer on ice for 10 min. The cell lysate was collected and added to 600 µL of MQ water before ultracentrifugation at 10,000 g at 10 °C for 30 mins. Next, the 500 µL was discarded and replaced with 500µL of MQ for a repeat round of ultracentrifugation under the same settings. Only the final 100 µL of solution was collected and processed for the proteomics study. For proteomics of the regurgitated clusters, HMC3 cells was exposed to BAS and Aβ for 24 hrs each and clusters were extracted by treating the cells with RIPA lysis buffer and clusters were purified from the cellular debris by washing with 100 kDa centrifugal filters in PBS. The purified clusters were processed for proteomics sample preparation. 300 µL of buffer (6M guanidine chloride, 50mM Tris pH 8 and 10mM DTT) was added to the digested fibril extract in a Lobind tube. The tube was then sonicated for 10 s in an ice water bath on Sonics Vibracell and vortexed at 30°C for 1 h. Next, 12.5µL of 25mM acrylamide was added to the sample and vortexed for another 1 h at 30°C. 2.6µL of 5mM DTT was transferred to each tube and frozen at −20°C overnight. The following day, the samples were centrifuged at room temperature, 18,000 g for 5 min and transferred to 10kDA cut-off Amicon column. Samples were further centrifuged at room temperature, 14,000 rpm for 40 min (setting I). 200 µL of 50mM ammonium bicarbonate was added to the sample and centrifuged with setting I. Finally, 130µL of 50mM ammonium bicarbonate and 1.5µL trypsin was used to incubate in a humid chamber at 37°C overnight. On the next day, the Amicon column was transferred to a new collection tube and centrifuged with setting I. Next, 50µL of 0.5M NaCl was added to the column and centrifuged again with setting I. The protein in the collection tube was collected in a ziptip (Merk, Australia) before being transferred to the School of Chemistry & Molecular Biosciences, Mass Spectrometry Facility, The University of Queensland for analysis.

### ASC speck formation in MDMi and THP1 cells

BAS (microbial seeds, concentration 0.1 µM). Aβ fibrils (1 µM). Methods for cell culture, differentiation, IL-1b and IL-6 measurement as described above Cell Culture section. As in Aβ + BAS samples, cells were exposed to BAS for 24 hrs, BAS media was removed and cells were exposed to Aβ for additional 24 hrs, we measured the ILs in BAS containing media that was removed prior to addition of Aβ.

### Cell stimulation with THP1 and MDMi cells

THP1-ASC-GFP, primary mouse microglia, and MDMi were seeded at 100,000 cells per well in 96-well plates or 500,000 cells per well in 12-well plates. For inflammasome activation, cells were treated with either BAS (0.1 µM) or Aβ fibrils (1 µM) alone for 24 hours or BAS for 24 hours then removed and treated with Aβ for extra 24 hrs, for NLRP3 inflammasome activation positive control cells were primed with 200 ng/mL Lipopolysaccharide (LPS; InvivoGen, #tlrl-3pelps) in serum-free medium for three hrs, followed by thorough washing to remove residual LPS and activated with Nigericin (10 µM) for 1 hours (**Fig. 4A**).

### ELISA and immunocytochemistry with THP1 and MDMi cells

Following incubation for the designated treatment periods, the supernatant was collected to quantify cytokine secretion. The concentration of IL-1β was measured using ELISA kits according to the manufacturer’s instructions Human IL-1β (R&D System, #DY201)

THP1-ASC-GFP cells and primary mouse microglia were seeded at 200,000 cells/well in poly-D-lysine-coated 12-well plates with a coverslip. THP1-ASC-GFP cells were differentiated into macrophages as previously described. For immunolabeling, cells were fixed with 4% PFA for 10 minutes, washed three times with DPBS, and incubated in blocking buffer (2% bovine serum albumin (BSA), 0.1% Triton-X (Merck. #9036-19-5) in DPBS) for one hour at room temperature.

Immunofluorescence imaging was performed using a Zeiss AxioScan Z1 Fluorescent Instrument. The number of positive cells or intensity-derived quantifications was analysed by the imaging software CellProfiler (version 4.2.1) and Fiji (Version: 2.1.0/1.53c), using the same pipeline for each sample in the same experiment. Custom Matlab R2018b (9.5.0.944444) scripts were developed to streamline high throughput imaging data.

### iPSCs derived organoid

Undirected human brain organoids were generated as previously described with minor modifications.^45^ Briefly, iPSC line A18945 was maintained on Matrigel (Corning) coated wells with mTeSr plus (Stemcell tech). Cells were dissociated with 2 minutes incubation with Accutase (Thermofisher) and followed by seeding of 12000 single cells in each well of a low attachment 96-well U-bottom low attachment plate (Corning) in mTeSr plus with 10 μM Y-27632 for two days to generate embryoid bodies (EBs). 80% of the medium was then replaced by EB medium for 6 days, followed by Neural induction media in the same 96-well plate. On day 11, EBs were transferred on sheets of parafilm, covered with 15 μl of 100% Matrigel (Corning) for embedding and incubated for 23 min at 37 °C. Embedded single EBs were then transferred to a 24-well plate in 1.5 ml of differentiation media per well, replaced every 3 days for 100 days. The microinjection of the organoids was performed on PV830 Pneumatic Picopump, WPI operating at 20 psi pressure using a pulled capillary tube needle with 100 nL injection volume per press. The agarose templates were prepared by adding 1% PenStrep in the agarose that was solidified under UV light at room conditions. During microinjection, the organoids were immersed in 1% PenStrep in DPBS. A total of 100 nL of the sample volume (BAS 0.1 µM with or without Aβ fibrils 1 µM, DPBS) were microinjected to the organoids. The microinjected organoids were maintained for 1 week in organoids medium and replaced every 2 days. On day 7, the organoids were fixed in 4% paraformaldehyde and sectioned into 16 µM thick sections on a cryostat. Organoids injected with prelabelled (AF) BAS and Aβ fibrils were used to study the colocalisation of BAS and Aβ with microglia. DAPI (1µg/mL) (Sigma # D9542), β3 tubulin and 1:500 (Sigma, # T8578) and Iba 1 1:500 (Wako, # 019-19741) staining was performed as previously described.^71^ Secondary antibodies conjugated to Alexa Fluor 488, 568, or 647 (Invitrogen) were employed for signal detection. Organoid sections were imaged under a Leica SP8 confocal microscope. The images were quantified for β3 tubulin integrity, fluorescence intensity and colocalization by Image J.

### Microinjection and oral gavage to adult zebrafish

The anesthetised fish was held on a Petri dish using two small pieces of foams (soaked with fish water) and held with forceps. An incision was made in the skull above cerebroventricular space using 1 mL syringe needle. Cerebroventricular injection of Aβ fibrils was performed with 1 µL Hamilton glass syringe and a total of 1 µL of the Aβ fibrils solution (100 µM in DPBS) was injected in between the right and left telencephalon with needle not penetrating more than 1 mm. Oral administration of BAS was performed by using a polypropylene feeding tube (Instech FTP-22-25) that was gently inserted no more than 15 mm into the fish for oral gavage. The oral gavage of BAS (10 µL) was performed in the adult fish with 1 µM of BAS (1:1 for FapC and CsgA) each day for three consecutive days and Aβ fibrils were injected to the brain on third day. After microinjection and oral gavage, the fish was placed back in the fish water tank to recover and placed back in the circulatory system. The zebrafish were grouped for samples of Aβ and BAS alone, BAS + Aβ and buffer control.

### Behaviour analysis and immunohistochemistry

The fish were maintained for 1 week and analysed for behavioural pathology. The swimming behaviour was recorded using zebrabox (viewpoint) in a 1 L swimming tank and the video (5 min observation) was analysed by Tox Trac software for the parameters of swimming trajectory, total distance travelled, % area explored and exploration rate (a.u.). The observations were made for 3 fish per sample and three times a day and then averaged together. For immunohistochemistry, the fish was euthanized in ice chilled water with 0.4% tricaine for 1 min and head of the fish was separated with a sharp blade at the pectoral fin. The head was fixed in 2.5% paraformaldehyde (PBS) for 24 hrs and then transferred to PBS and stored. The head was dehydrated by placing in gradually increasing concentration of ethanol (20-100%) and then placed in 20% sucrose overnight. The head was mounted with Tissue Trek OCT medium at −20 °C and sectioned into 20 µM thick sections with a cryostat. The sections were mounted on glass slides and process for immunostaining. The tissue sections were washed with DPBS thrice to remove OCT media, placed in antigen retrieval buffer (1M Sodium Citrate, 0.25% SDS, pH 6.0) and exposed to microwave for 5 min using PELCO BioWave microwave. The sections were washed with PBS and encircles with hydrophobic PapPen. The sections were rinsed with blocking buffer (5% BSA, 0.05% saponin, 0.05% sodium azide in PBS) and blocking buffer was used to dilute primary and secondary antibodies. The primary antibodies of Aβ (mouse monoclonal, Anaspec, AS-55922, 1:500 dilution), synaptophysin (Abcam, ab32594, 1:500 dilution) and Iba1 (GeneTex, GT10312, 1:500 dilution) were incubated with the tissue sections at 4 °C in a humidified chamber overnight. The secondary antibodies (1:1000 dilutions) were incubated with the tissue sections for 4 hrs at room temperature in a humidified chamber. For DAPI staining, Hoechst (1 µg/mL) was incubated with the tissue sections for 10 mins and washed with PBS. The washed sections were mounted with cover slip using DAPCO mounting medium. TUNEL assay was performed with Click-iT Plus kit (Invitrogen C10619) as per manufacturer directions. The slides were imaged under a fluorescence microscope and fluorescence intensities were measured by Fiji ImageJ software analysis.

### Statistics

All the experiments in the manuscript were repeated three times, unless otherwise specified. Data presented as mean ± the standard error of the mean (SEM). Students t test, one sample t and Wilcoxon test using GraphPad Prism 9.4 software was used for significance. P values more than 0.05 were considered insignificant.

## Acknowledgements

This work was supported by NHMRC Investigator grants (APP2009991 Javed, APP1197373 Davis and APP2009957 Woodruff). TEM was performed at the Centre of Microscopy and Microanalysis at UQ, experiments with juvenile zebrafish were performed at UQ aquatic facility and confocal imaging was performed at Australian National Fabrication Facility – Queensland Node.

## Author Contributions

I.J., H.F. and T.P.D., conceived the project. T.P.D. and I.J., arranged the funding. H.F. performed imaging, image analysis, western blotting, live cell imaging. E.A.B., J.A. and T.M.W performed the MDMi and THP1 culture, ASC imaging and cytokine measurement. G.P. and E.J.W conducted organoids experiments and immunohistochemistry. J.Z. and R.Q. performed HMC3 culture and J.Z. prepared the figures. K.H.K.C and M.S.T conducted proteomics and analysed the data. S.A.A measured IL from organoids and zebrafish immunohistochemistry. A.K. performed TEM imaging and graphical illustrations. I.J performed the microinjections and zebrafish experiments. D.E.O. synthesised FapC, CsgA proteins by recombinant expression and prepared amyloids and BAS seeds. I.J., H.F. and T.P.D. wrote the manuscript and E.A.B., J.A., G.P., T.M.W. and D.E.O edited the manuscript. All authors agreed on the presentation of the paper.

