## Supplementary data for "Interaction of Gut-Microbial Amyloids with Endogenous Amyloids can Drives Microglial Hyperactivity and Neuroinflammation in Alzheimer’s Disease"

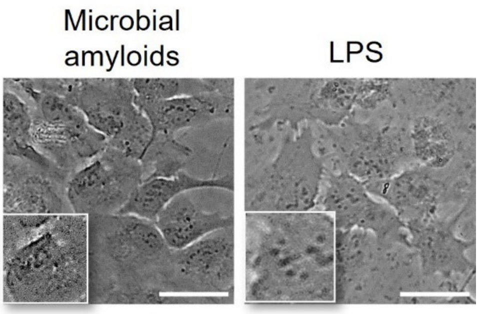


**Figure S1.** HMC3 microglia cells exposed to (left) full length microbial amyloids without Aβ or (right) lipopolysaccharides (LPS) as positive control did not develop intracellular vesicular aggregates (scale bar 20 μm).

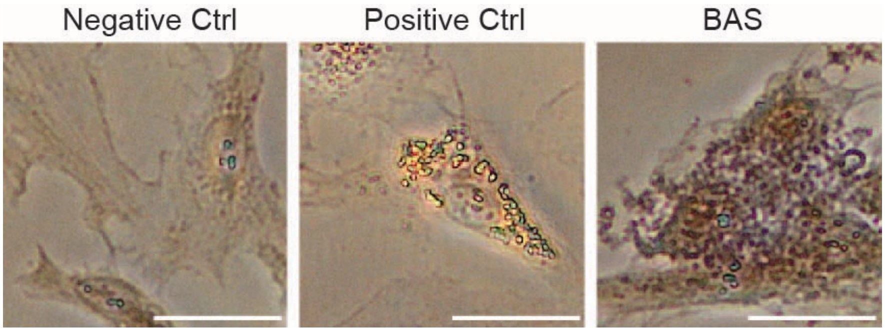


**Figure S2.** Exposure to BAS resulted in formation of intracellular aggregates that were negative for Oil RedO staining for lipid droplet formation (scale bar 50 μm). Untreated cells were used as negative control while. The brown color in BAS panel represents intracellular granules formed in microglia upon BAS exposure.

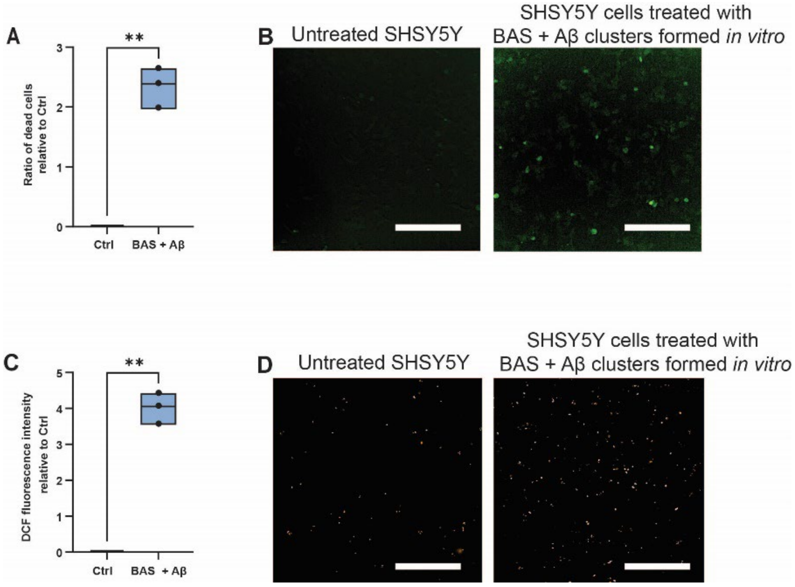


**Figure S3.** BAS were physically mixed with Aβ fibrils to assemble into clusters. These *in vitro* formed clusters were exposed to SHSY5Y cells that resulted in higher cytotoxicity (A, B) and ROS (C, D) generation in the cells- This is consistent with the increased cytotoxicity and ROS observed when SHSY5Y cells were co-cultured with BAS + Aβ exposed microglia (**Fig. 2E, F**). Scale bar in B, D 500 µm.

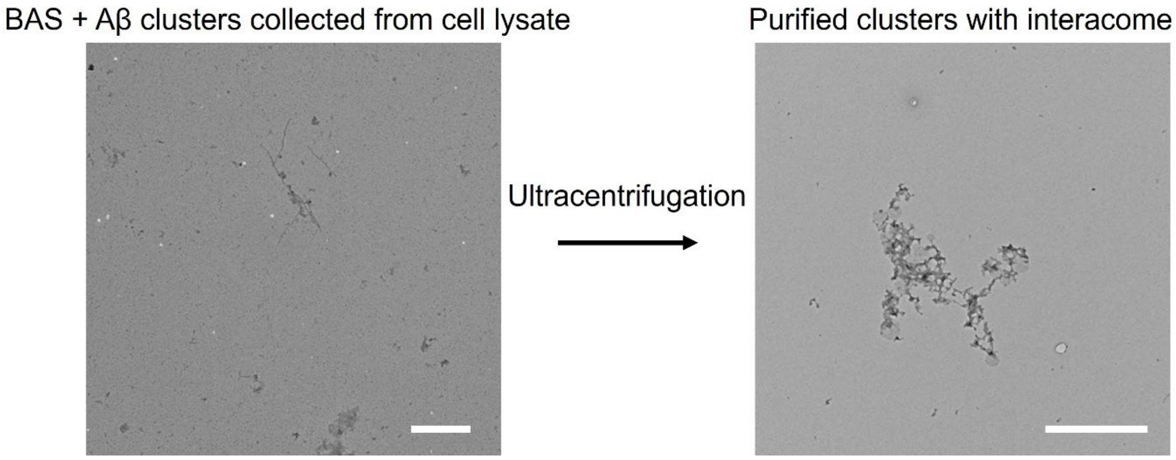


**Figure S4.** HMC3 microglia cells incubated with BAS + Aβ were lysed, and lysate was collected and purified by repeated ultracentrifugation. The clusters, together with their interactome developed inside the cells were subjected to proteomics analysis.

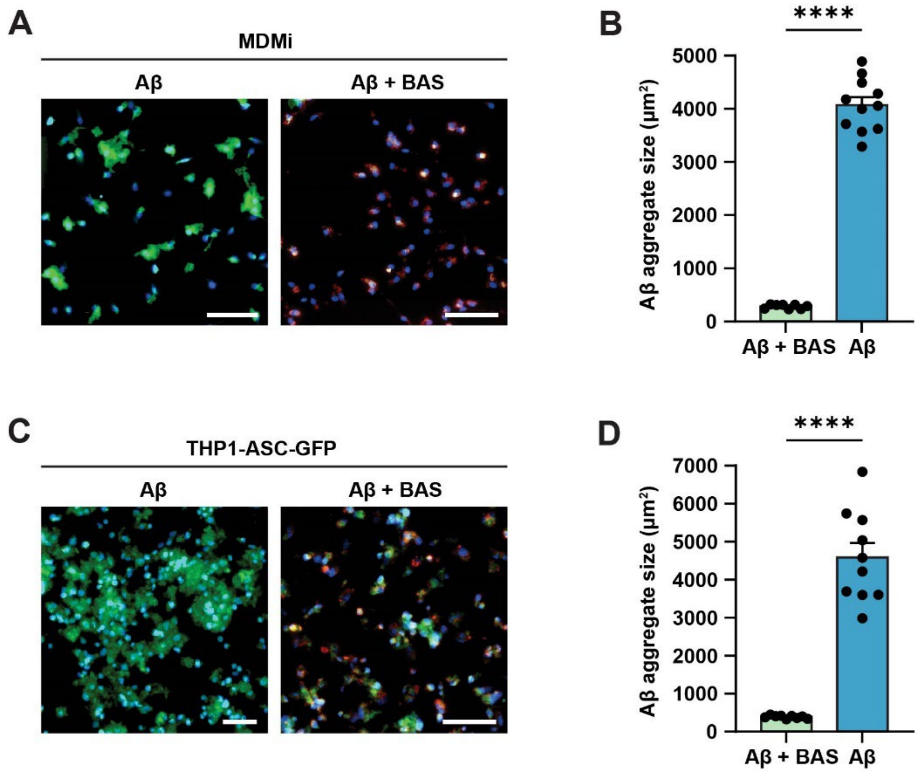


**Figure S5.** In MDMi and THP1 cells, subsequent phagocytosis of Aβ fibrils after BAS, resulted in compaction of the size of Aβ into smaller aggregates. The fluorescent images are presented in A and C for MDMi and THP1 cells (scale bar 200 μm) while size of the Aβ plaques quantified and presented in graphed data in B and D. Aβ fibrils alone, when exposed to MDMi or THP1 cells alone, resulted in the formation/sedimentation of fluffy larger sized plaques on the cells.
